# Single cell data enables dissecting cell types present in bulk transcriptome data

**DOI:** 10.1101/2024.07.30.605755

**Authors:** Wasco Wruck, James Adjaye

## Abstract

The quality of organoid models can be assessed by single-cell-RNA-sequencing (scRNA-seq) but often only bulk transcriptome data is available. Here we present a pipeline for the analysis of scRNA-seq data and subsequent “deconvolution”, which is a method for estimating cell type fractions in bulk transcriptome data based on expression profiles and cell types found in scRNA-seq data derived from biopsies. We applied this pipeline on bulk iPSC-derived kidney and brain organoid transcriptome data to identify cell types employing two scRNA-seq kidney datasets and one brain dataset. Relevant cells present in kidney (eg. proximal tubules, distal convoluted tubules and podocytes) and brain (eg. neurons, astrocytes, oligodendrocytes and microglia) with obligatory endothelial and immune-related cells were identified. We anticipate that this pipeline will also enable estimation of cell type fractions in organoids of other tissues.

## Introduction

Organoids are three-dimensional (3D) tissue cultures resembling organs. They contain multiple cell types with the ultimate goal to mimic all cell types in the same fractions and organization of the real organ. A plethora of already existing and envisioned applications range from developmental and disease models to regenerative medicine and transplantable organs. An important step needed for assessing the quality of iPSC-derived organoids is to determine the fractions of cell types present. Single-cell RNA-seq (scRNA-seq) techniques can be employed to calculate these cell type fractions in high accuracy. However, as scRNA-seq techniques are very expensive not every lab can afford to use this approach. An accurate estimation of cell type proportions in organoids from bulk transcriptome data therefore would be of great benefit to enable cell type estimations from existing bulk trancriptome data or for projects without the necessary funding for scRNA-seq experiments.

The so-called “deconvolution” methods are based on the idea that in the bulk transcriptome, gene expression profiles are merged, “convolved” by a function which can be determined and reverted with the abundant scRNA-seq information. By application of the reverse-the “deconvolution” function, one can dissect bulk transcriptome data into expression profiles and proportions of the contained cell types. Several “deconvolution” tools have been introduced including CIBERSORTx^7^, MuSiC ^2^, BisqueRNA ^3^, BayesPrism ^4^ and InstaPrism ^5^. While in terms of accuracy BayesPrism ^4^ out-performs the other approaches, InstaPrism ^5^ re-implements BayesPrism in a more memory- and runtime-efficient fashion. For the analysis of scRNA-seq data the R package Seurat ^6^ provides a sophisticated workflow with implementations of all required methods. Seurat can identify clusters of similar cells in scRNA-seq data and identify markers characteristic of these cell clusters. Cell types can be associated with clusters via these markers. While clusters can be annotated manually with cell-types based on a priori known canonical markers, the R package scMayoMap ^7^ offers a semi-automated approach for cell-type-cluster association based on a huge database of cell-type-specific expression profiles. Deconvolution tools have been applied for the estimation of cell types in tumors ^4, 8^ or in tissue biopsies ^9^ or other biological samples ^10^. However, studies investigating cell type deconvolution in organoids are scarce. To our knowledge there is a study using deconvolution via CIBERSORTx in colon cancer organoids ^11^ and the tool VoxHunt using deconvolution for prediction of organoid brain regions based on the Allen Brain Atlas which although very useful is restricted to these brain datasets ^12^.

Here, we introduce a pipeline for deconvolution of bulk organoid transcriptome data using scRNA-seq datasets of various tissues and demonstrate its performance with use cases of bulk transcriptome data from kidney ^13^ and brain organoids ^22, 21^ employing publicly available (NCBI GEO) scRNA-seq kidney datasets GSE241302 and GSE202109 associated with studies by Shi et al. ^14^ and McEvoy et al. ^15^ and brain datasets GSE104276 associated with the study by Zhong et al. ^16^ .

## Results

### A pipeline for the prediction of cell type proportions in bulk transcriptome data via scRNA-seq data

Figure 1 shows the pipeline for the deconvolution of bulk transcriptome data to single cell types using single-cell sequencing (scRNA-seq) data. ScRNA-seq data are analyzed via the R package Seurat ^6^ . Crucial steps here include quality control to filter out cells of bad quality based on thresholds for mitochondrial content and number of expressed features. However, these thresholds need fine-tuning to account for characteristics of the cell types, e.g. high mitochondrial density in kidney cells. Clusters of distinct cell types are found by the Seurat routine *FindClusters* but requiring appropriate adjustment of the resolution parameter to detect all required cell types. Marker genes are identified by the routine *FindAllMarkers*. Cell types are assigned to the single-cell clusters using the predictions from the scMayoMap package ^7^ . Predictions are checked and curated manually, e.g. for missing cell type annotations employing the cell type enrichment analysis from the EnrichR webtool ^17^ . Bulk transcriptome data from RNA-seq or microarray experiments and scRNA-seq data are merged using a reduced geneset of several thousand markers found via Seurat. scRNA-seq and RNA-seq data are brought to the same level with the *ComBat* batch effect removal method from the R package sva ^18^. Resulting data are normalized with quantile normalization ^19^ . The normalized data are used as input to the R package InstaPrism to predict proportions of cell types prevalent in the bulk data. InstaPrism is a fast implementation of BayesPrism which employs a Bayesian approach to estimate gene expression in single cell types based on the sc- and bulk-RNA-seq data.

**Figure 1:**
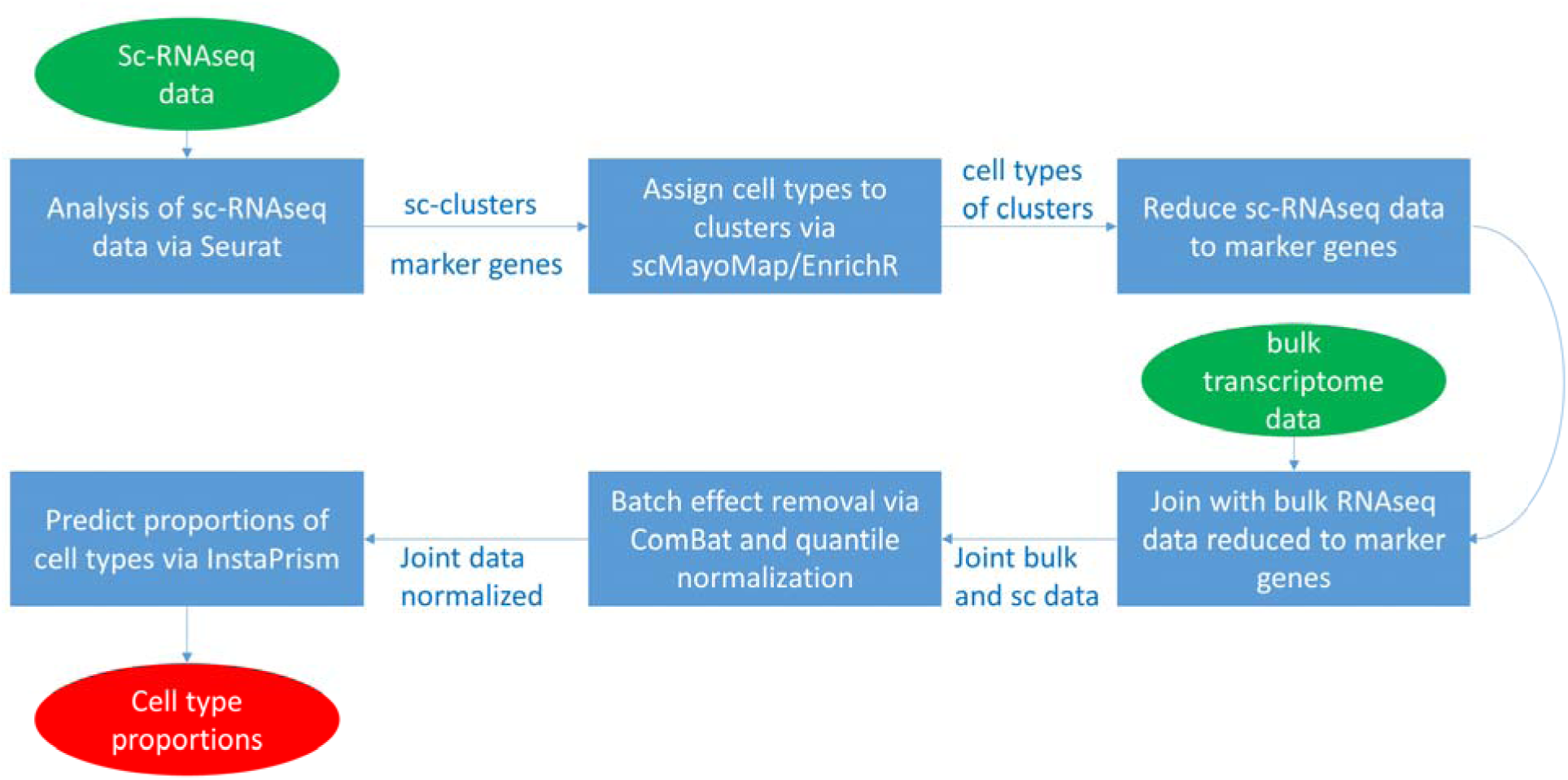
Bulk transcriptome data can be deconvoluted to single cell types using single-cell sequencing data as reference. ScRNA-seq data are analyzed via the R package Seurat to identify single-cell clusters and marker genes. Cell types are assigned to the single-cell clusters by cell type enrichment analysis with the scMayoMap package ^7^ and the EnrichR ^17^ webtool. The scRNA-seq data is reduced to the several thousand marker genes found by Seurat. Bulk transcriptome data from RNA-seq or microarray experiments are joined with the scRNA-seq data. Batch effects are removed with the ComBat method from the R package sva and data are normalized with quantile normalization. Proportions of cell types prevalent in the bulk data are predicted via the R package InstaPrism which is a fast implementation of BayesPrism. The R package uses a Bayesian approach to estimate gene expression in single cell types based on the sc-and bulk-RNA-seq data.

### Cluster analysis of scRNA-seq data and cell-type assignment

The granularity of clusters resulting from the scRNA-seq analysis depends on fine-tuning of parameters such as the resolution in the Seurat software. If the resolution is too small, cell types with lower abundance such as podocytes may not be identified. We set the resolution to 0.8 to find the clusters displayed in figure 2a including podocytes. A disadvantage of a too high resolution is that cell types may be spread over several clusters - such as the proximal tubules (PT) in this case - which on the other hand may reflect subtypes. However, podocytes are detected only with a high resolution. Cell types are assigned to the clusters found by Seurat using a comprehensive collection of scRNA-seq and RNA-seq datasets provided by the R package scMayoBase. This revealed distinct kidney cell types including podocytes, PTs, DCTs and also immune-system-related cells (Figure 2b). Clusters 13 and 24 yielded no result in scMayoBase and were annotated via the EnrichR webtool (using ARCHS4 tissues datasets) as dendritic cells (Table S2, EnrichR annotations). Fractions are calculated by summing up cell types associated with single cells (Table 1). PT is the most abundant cell type with about 42%, followed by LOH thin descending limb (LOH_thin) with about 13%, intercalated cells about 9%, LOH thick ascending limb (LOH_TAL) about 8%, endothelial cells with about 5%, B-cells with about 4%, T-cells, dendritic cells and fibroblasts each about 3%, principal cells and macrophages each about 2%, Natural Killer (NK) cells about 1.5%, Distal convoluted tubule (DCT) cells about 1.3%, neutrophils about 1.2%, erythroid precursor cells about 0.8% and podocytes about 0.2%.

**Figure 2:**
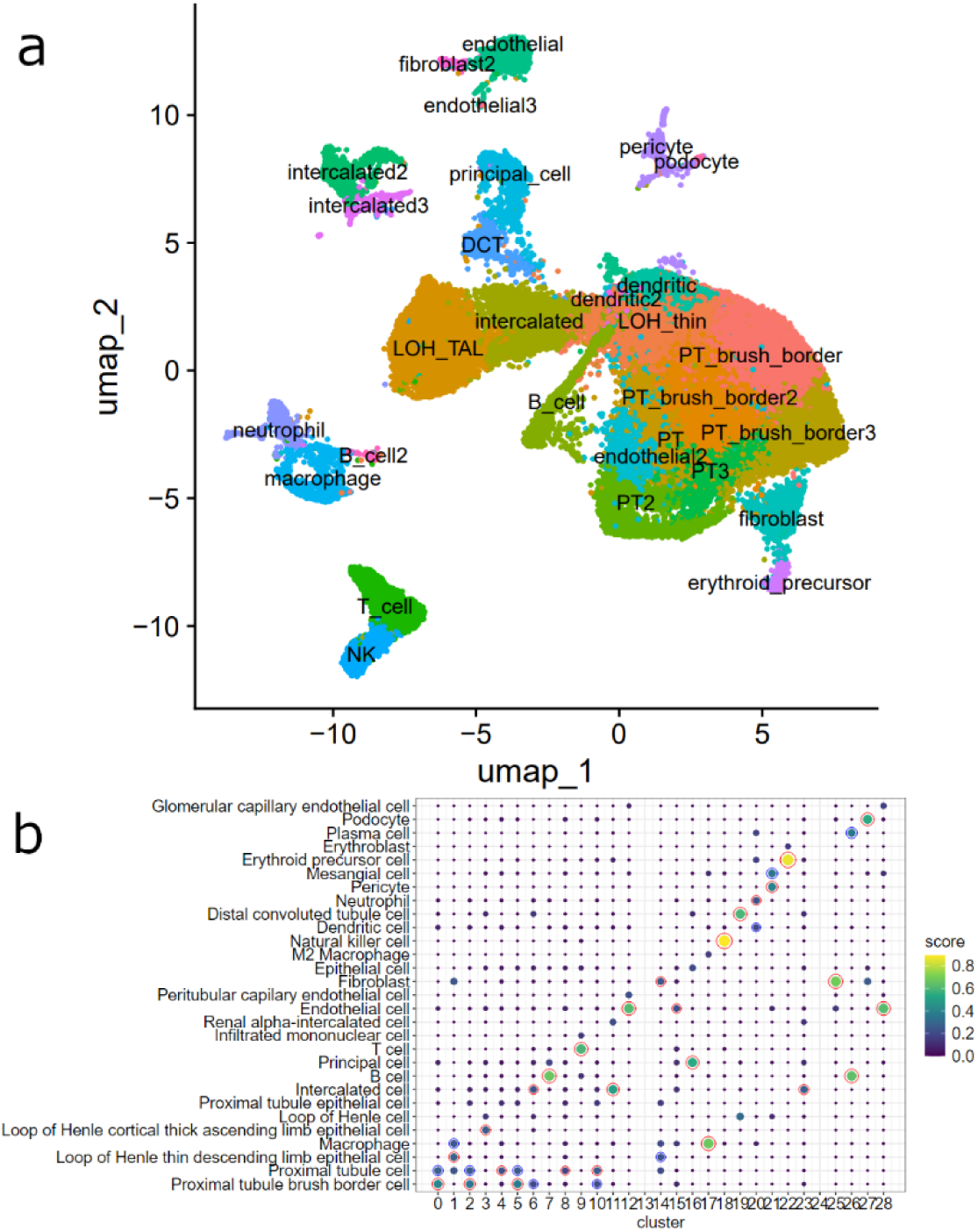
Distinct kidney and immune cell types are identified via cluster analysis and assignment via scMayoMap. (a) UMAP plot of clusters found via Seurat in the scRNA-seq kidney dataset GSE241302. (b) Cell types were assigned to clusters via scMayoMap yielding most relevant kidney cell types and also cell types associated with the immune system. Red circles highlight the top scores, blue circles 2^nd^ top scores. Dot colour and size reflect the score.

**Table 1:**
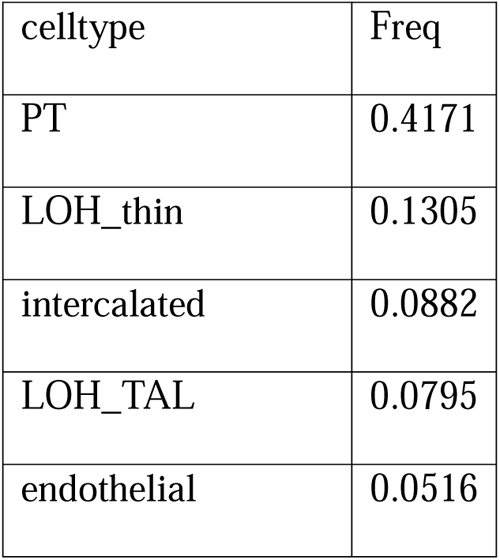
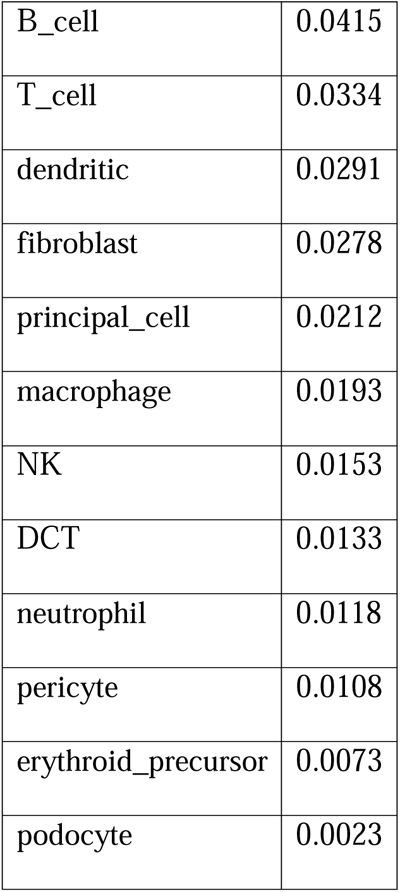
Fractions of celltypes in scRNA-seq kidney dataset GSE241302. PT: Proximal tubule, DCT: Distal convoluted tubule, LOH: Loop of Henle, NK: Natural-killer-cells.

### Markers of distinct kidney and immune cell types are expressed in iPSC-derived kidney organoids

A result of the Seurat analysis is a table of markers of the identified clusters (table S3). We used the 50 most significant markers (by Seurat p-value) for relevant kidney and immune-related cell types to assess their expression in the kidney organoids from our previous publication-Nguyen et al. ^13^ . The expression of the marker genes in kidney organoids with and without PAN treatment is shown in proximal tubule cells including marker genes CUBN and LRP2 (Figure 3a), (Figure 3b) Loop of Henle (LOH) cells including marker gene UMOD, (Figure 3c) distal convoluted tubule (DCT) cells including marker gene SLC12A3, (Figure 3d) podocytes including marker genes NPHS2 and PODXL, (Figure 3e) T-cells including marker gene CD3D, (Figure 3f) endothelial cells including marker genes PECAM1 (CD31), EMCN and FLT1, (Figure 3g) macrophages including marker genes CD68 and CCL3, (Figure 3h) pericytes including marker gene PDGFRB.

**Figure 3:**
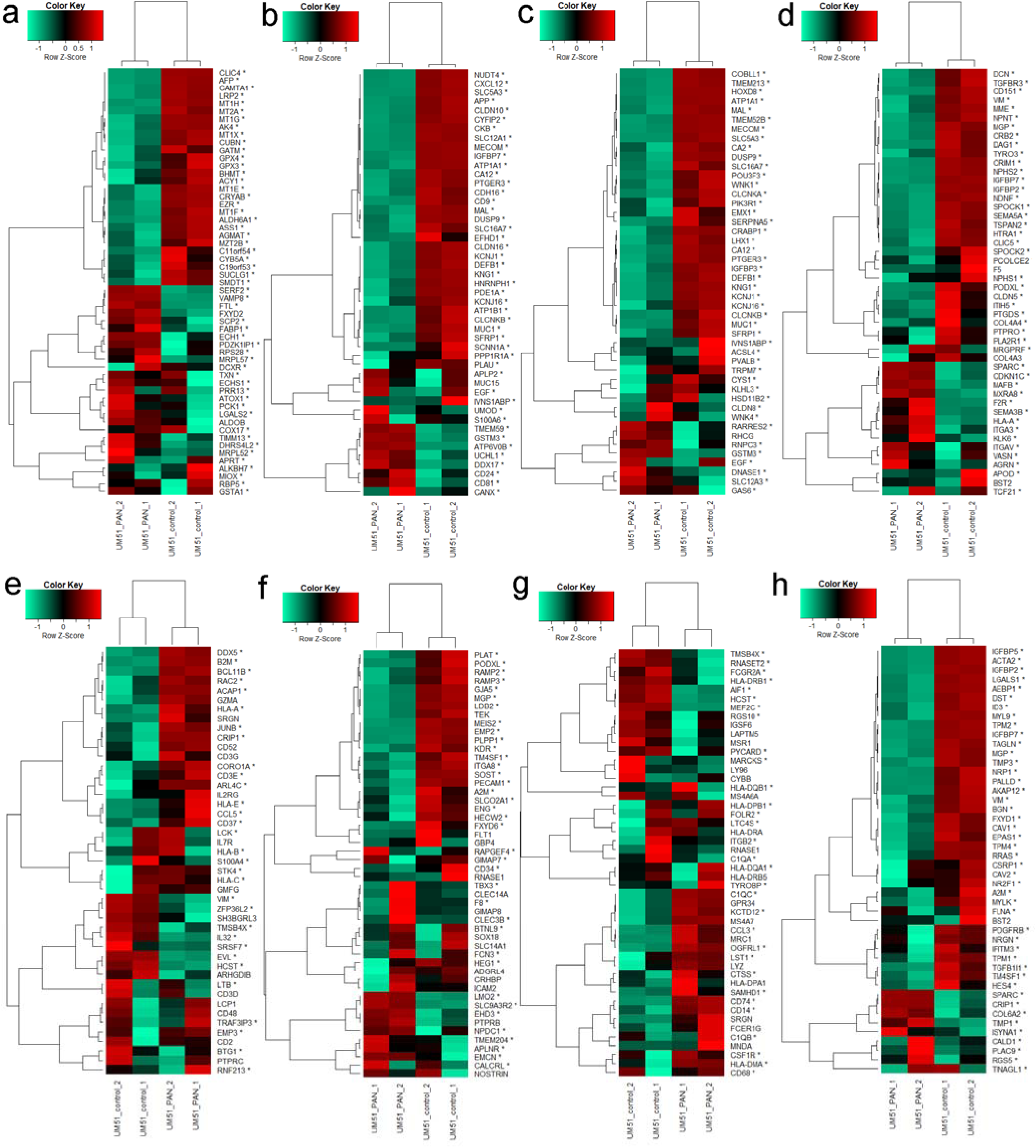
Markers of distinct kidney, endothelial, pericyte and immune cell types are expressed in iPSC-derived kidney organoids. The heatmaps show the expression in kidney organoids (with and without PAN treatment and iPSCs cultured as spheroids) of the 50 most significant (by Seurat marker p-value) marker genes for each cell type. (a) proximal tubule cells including marker genes-CUBN and LRP2 (64 genes), (b) Loop of Henle (LOH) cells - UMOD, (c) distal convoluted tubule (DCT) cells-SLC12A3, (d) podocytes-NPHS2 and PODXL, (e) T-cells-CD3D, (f) endothelial cells-PECAM1 (CD31), EMCN and FLT1, (g) macrophages-CD68 and CCL3, (h) pericytes-PDGFRB.

### Deconvolution of bulk kidney organoid RNA-seq data

We set out to investigate cell types in bulk microarray transcriptome of kidney organoids from our previous publication-Nguyen et al. ^13^ using the results of the scRNA-seq kidney data analysis. As illustrated in the pipeline scheme in Figure 1, we applied the InstaPrism deconvolution to the bulk kidney organoid transcriptome data normalized together with the scRNA-seq kidney data from GSE241302. The pie chart in Figure 4a depicts the proportions of cell types identified in the untreated kidney organoids. Most abundant are proximal tubule cells (PT, PT_brush_border, 35%), followed by intercalated cells (11%), dendritic cells (10%), endothelial cells (7%), fibroblasts (6%), Loop of Henle thin descending limb (LOH_thin, 5%), Loop of Henle thick ascending limb (LOH_TAL, 4%), distal convoluted tubule cells (DCT, 4%), erythroid precursor cells (4%), B-cells (3%), pericytes (2%), principal cells (2%), macrophages (1.9%), podocytes (1.7%), T-cells (1.4%) natural killer cells (NK, 1.0%), and neutrophils (1.0%). The most abundant cell types are distributed over multiple sub-clusters potentially reflecting sub-cell-types. Figure 4b additionally shows the summarized proportions of kidney organoids treated with the nephrotoxin-PAN and the highest difference in proximal tubule cells, possibly reflecting tubular damage by PAN treatment as reported in our previous publication-Nguyen et al..

**Figure 4:**
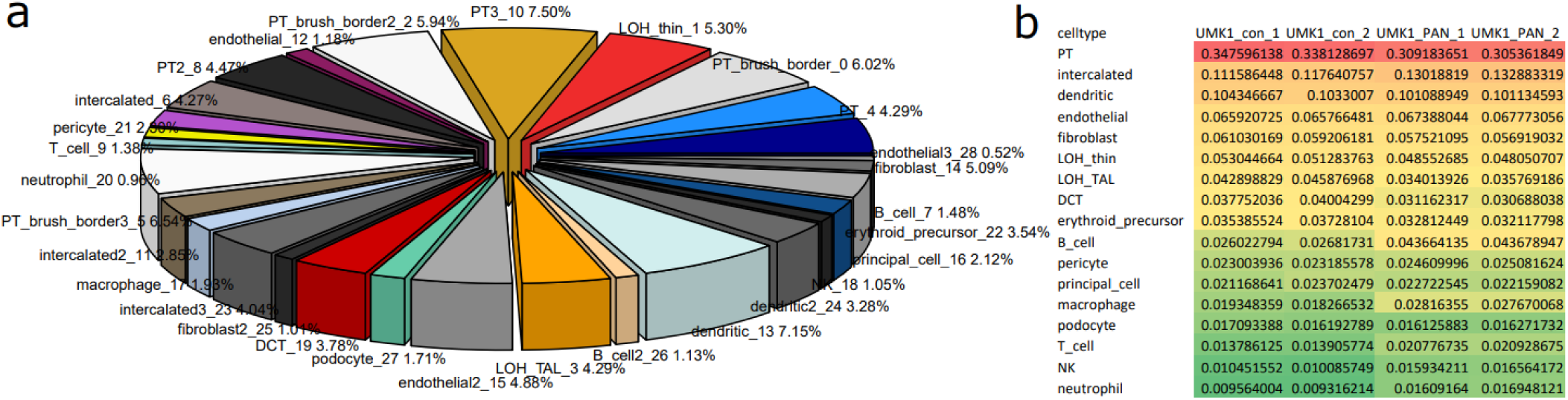
Cell type predictions via InstaPrism using the single-cell dataset GSE241302 for iPSC-derived kidney organoids. (a) Proportions of cell types predicted by InstaPrism for the bulk transcriptome samples of untreated kidney organoids. PT and DCT cells are the most abundant cell types in the untreated kidney organoids. (b) Proportions of cell types in bulk kidney organoid transcriptomes with and without PAN treatment.

### Deconvolution with another scRNA-seq dataset (GSE202109)

We now wanted to investigate the reproducibility of the predicted cell types in bulk microarray transcriptome of our kidney organoids using the scRNA-seq kidney dataset GSE202109. First, we analyzed the scRNA-seq data with Seurat and annotated clusters with scMayoMap. The UMAP (Uniform Manifold Approximation and Projection) plot in Figure 5a and the dot plot in Figure 5b show that essentially all relevant kidney cell types were identified. However, Table 2 shows that there was a very high abundance of 84% proximal tubule cells (PT, PT_brush_broder) while all other cell types were below 5%. This is propagated to the deconvolution and reflected in the pie chart in Figure 5c depicting the proportions of cell types found via InstaPrism deconvolution in our untreated kidney organoids. Most abundant are proximal tubule cells (PT, PT_brush_border, between 66-67%), and amongst the other celltypes intercalated with about 8%, mesangial and DCT with about 4%, dendritic with about 3%, LOH_TAL with about 2.5%, principal cells with about 2%, NK cells with about 1.9%, podocytes with about 1.8%, pericytes with about 1.6%, endothelial cells with about 1.2% and fibroblasts with about 1.1%.

**Figure 5:**
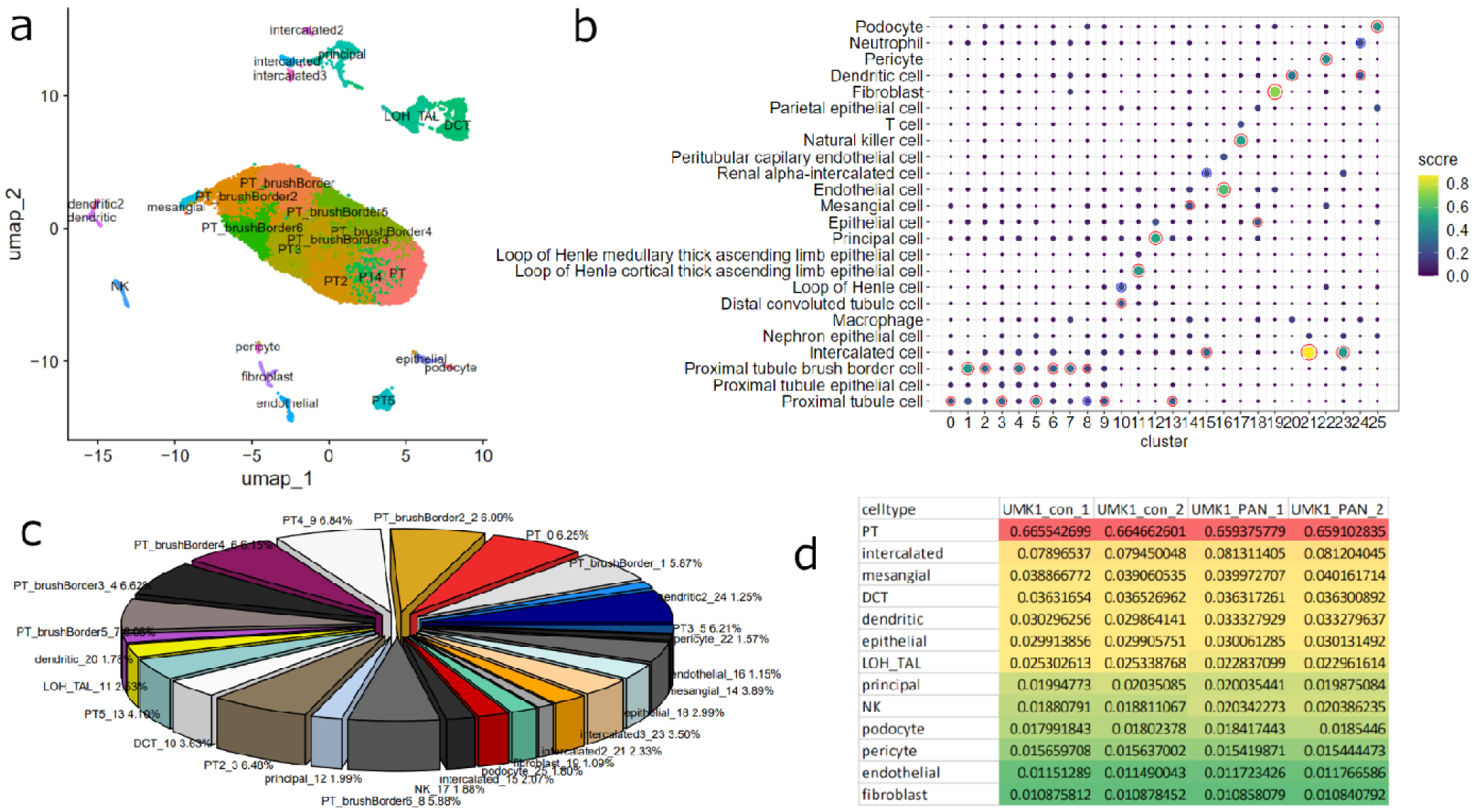
Distinct kidney and immune cell types are also found via scRNA-seq dataset GSE202109. (a) UMAP plot of clusters found via Seurat in the scRNA-seq kidney dataset GSE202109. (b) Cell types were assigned to clusters via scMayoMap yielding most relevant kidneycell types and also cell types associated with the immune system. Red circles highlight the top scores, blue circles 2nd top scores. Dot color and size reflect the score. (c) Proportions of cell types predicted by InstaPrism for the bulk transcriptome samples of untreated kidney organoids using scRNA-seq kidney dataset GSE202109. PT cells are the most abundant cell types in the untreated kidney organoids. (d) Proportions of cell types in bulk kidney organoid transcriptomes with and without PAN treatment (using GSE202109).

**Table 2:** Fractions of celltypes in scRNA-seq kidney dataset GSE202109. PT: Proximal tubule, DCT: Distal convoluted tubule, LOH: Loop of Henle, TAL: Thick ascending limb, NK: Natural-killer-cells.

| celltype | Freq |
| --- | --- |
| PT | 0.8393 |
| DCT | 0.0460 |
| LOH_TAL | 0.0259 |
| principal | 0.0216 |
| intercalated | 0.0151 |
| mesangial | 0.0105 |
| endothelial | 0.0088 |
| NK | 0.0086 |
| epithelial | 0.0082 |
| dendritic | 0.0059 |
| fibroblast | 0.0052 |
| pericyte | 0.0030 |
| podocyte | 0.0017 |

### Deconvolution of bulk brain organoid data with the developmental prefrontal cortex scRNA-seq dataset GSE104276

Moreover, we wanted to investigate the prediction of cell types in bulk microarray transcriptome of brain organoids using the scRNA-seq prefrontal cortex (PFC) dataset GSE104276. We analyzed the scRNA-seq data with Seurat and annotated clusters with scMayoMap. The UMAP plot in Figure 6a and the dot plot in Figure 6b show the results and detected relevant brain cell types such as neurons, astrocytes, oligodendrocytes, microglia, interneurons and precursor cells of neurons and oligodendrocytes. We annotated cluster 1 with interneurons having the highest score and GABAergic neurons the second highest as “GABAergic neurons” to be more specific and also include inhibitory neuron types as other author have found a distribution of 55% Excitatory, 35% non-neuron and 11% Inhibitory neurons in mouse PFC ^20^. Table 3 shows that neurons of distinct types have the highest abundances in our human data while there are also considerable proportions of 5-6% astrocytes and oligodendrocytes each and 3% microglia. Additionally, precursor cells such as oligodendrocyte precursor cells (3%) and neural stem cells (11%) were detected in the scRNA-seq data. The results from InstaPrism-based deconvolution of bulk organoid transcriptome data from our previous publication-Pranty et al. ^21^ is presented as a pie chart in Figure 6c and depicts the proportions of cell types found in untreated brain organoids, the most abundant are Interneurons (15.2%). In another earlier study from our group, we developed a brain organoid model to study the molecular mechanisms associated with Nijmegen-Breakage syndrome (NBS) ^22^ . Results of the cell type predictions via InstaPrism for the NBS brain organoid model based on the same scRNA-seq dataset GSE104276 are shown in Figure 6e and f. The overall distribution of cell types in both brain organoid models is similar. Most prominent is the difference in microglia which are more abundant in the control UJ samples in the model by Pranty et al. (Figure 6c, 5.65%) than in the one by Martins et al. (Figure 6e, 0.51%). However, in the second control (A4) in the brain organoid model by Martins et al. microglia have a similar abundance (Figure 6f, 6%) as in the model by Pranty et al. pointing at individual differences between organoids derived from distinct iPSC lines.

**Figure 6:**
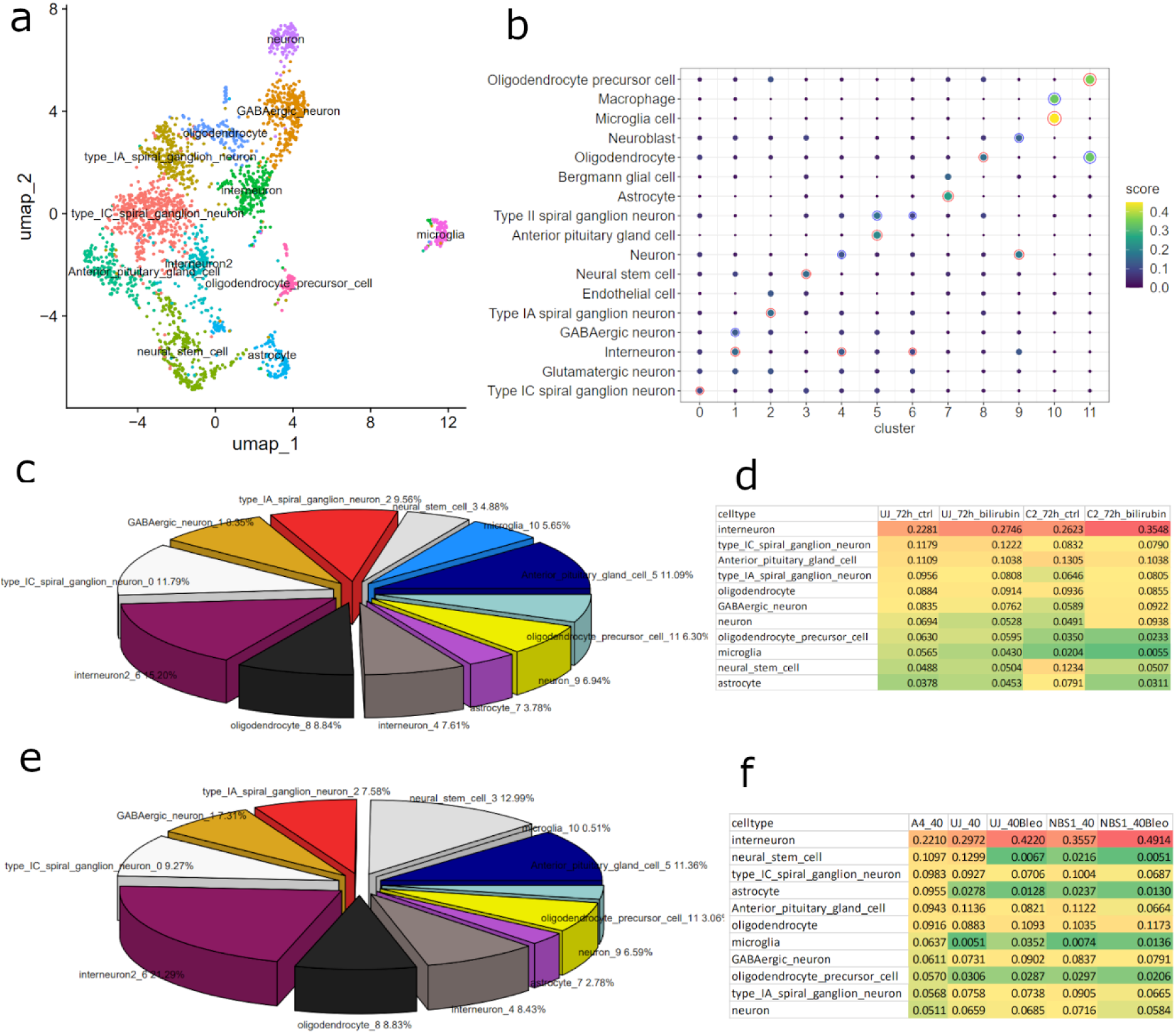
Distinct brain cell types are found in bulk brain organoids via the developmental prefrontal cortex (PFC) scRNA-seq dataset GSE104276. (a) UMAP plot of clusters found via Seurat in the scRNA-seq PFC (from gestational weeks 8 to 26) dataset GSE104276. (b) Cell types were assigned to clusters via scMayoMap to unveil brain cell types. Red circles highlight the top scores, blue circles 2nd top scores. Dot colour and size reflect the score. (c) Proportions of cell types predicted by InstaPrism for the bulk transcriptome sample UJ of untreated brain organoids from our previous publication (Pranty et al. ^21^) using scRNA-seq PFC dataset GSE104276. Inter-neurons are the most abundant cell types in the untreated brain organoids. (d) Proportions of cell types in bulk brain organoid transcriptomes (Pranty et al. ^21^) with and without bilirubin treatment (UJ: brain organoids derived from control iPSCs, C2: brain organoids derived from iPSCs with Crigler-Najar mutation in UGT1A1). (e) Proportions of cell types predicted by InstaPrism for the bulk transcriptome sample UJ of untreated brain organoids from our previous publication (Martins et al. ^22^) using scRNA-seq PFC dataset GSE104276. Interneurons are the most abundant cell types in the untreated brain organoids. (f) Proportions of cell types in bulk brain organoid transcriptomes (Martins et al. ^22^) with and without Bleomycin treatment (UJ and A4: brain organoids derived from control iPSCs, NBS1: brain organoids derived from iPSCs with Nijmegen-Breakage-Syndrome mutation in NBN).

**Table 3:**
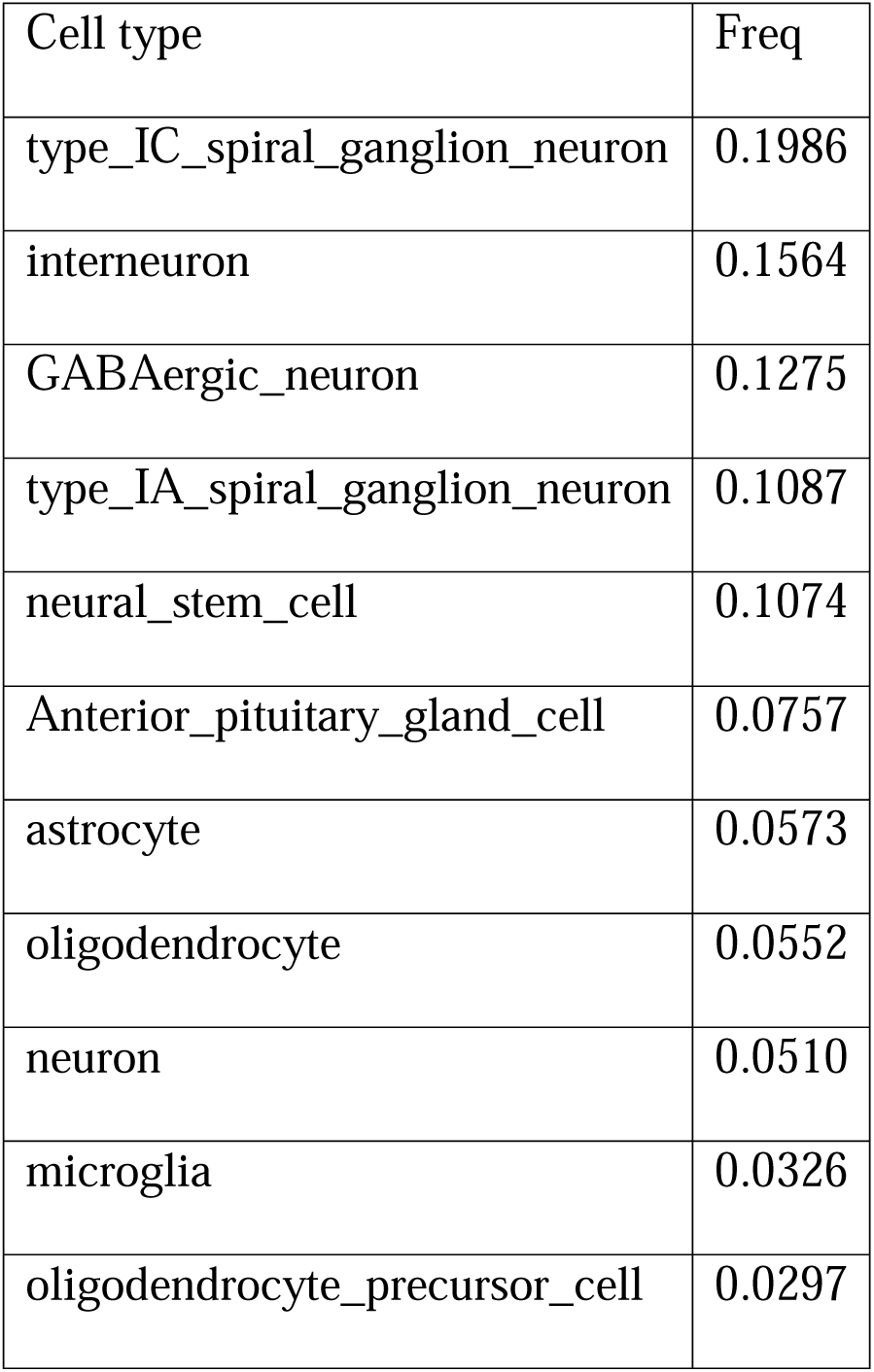
Fractions of celltypes in scRNA-seq prefrontal cortex dataset GSE104276.

| Cell type | Freq |
| --- | --- |
| type_IC_spiral_ganglion_neuron | 0.1986 |
| interneuron | 0.1564 |
| GABAergic_neuron | 0.1275 |
| type_IA_spiral_ganglion_neuron | 0.1087 |
| neural_stem_cell | 0.1074 |
| Anterior_pituitary_gland_cell | 0.0757 |
| astrocyte | 0.0573 |
| oligodendrocyte | 0.0552 |
| neuron | 0.0510 |
| microglia | 0.0326 |
| oligodendrocyte_precursor_cell | 0.0297 |

## Discussion

The aim of this study was to analyze the distribution of cell types in bulk transcriptome data of iPSC-derived kidney and brain organoids using the cell-type-specific gene expression profiles provided by biopsy-based kidney and brain scRNA-seq data. For this task we assembled a pipeline including the main processes of cluster analysis of single-cell data, cluster annotation with cell types based on the comprehensive databases of scMayoBase and deconvolution of bulk data via the R package InstaPrism to estimate the cell type distributions. We tested our pipeline with scRNA-seq data from kidney biopsies and developmental brain and bulk transcriptome datasets of kidney and brain organoids. The results of the cluster analysis and annotation of the two kidney single-cell datasets showed that all relevant kidney cell types were identified. However, there were differences in the details and in the distribution of cell types. Most prominent was the difference between about 42% PT (proximal tubule) cells in dataset GSE241302 and about 84% PT cells in dataset GSE202109. This may be due to differences in the sample collection which were biopsies of the idiopathic membranous nephropathy patients and excision of paracancerous tissue (2 cm away from the tumor) of the control patient for dataset GSE241302 ^14^ and pre-implantation biopsies from living donor kidneys for dataset GSE202109^15^. Furthermore, there were differences in the extraction protocols as described in the associated publications by Shi et al. ^14^ and McEvoy et al. ^15^. Another single-cell transcriptomics study by Hinze and colleagues ^23^ also identified PT as the most abundant cell type in kidneys of healthy controls and AKI (acute kidney injury) patients but on a level more similar to the 42% PT we found in GSE241302. Thus, we suggest that the proportions of dataset GSE241302 are more representative for an average kidney. Finding a dataset which is representative for a standard average healthy human kidney is a challenge which will be influenced by several factors such as cortex/medulla, male/female, age, disease. These influences may be tackled by generation of multiple distinct reference datasets, e.g. for medulla and cortex, both sexes and distinct age groups. Nevertheless, a dataset accurately reflecting the composition of kidney cell types is of crucial relevance for the estimation of cell types in the bulk organoids. Another source of variation is the annotation which often is performed using one or two known marker genes. However, these are prone to bias and thus we decided to annotate via the scMayoBase tool. The authors of scMayoBase have demonstrated the superior accuracy and performance of their tool compared to other existing annotation methods^11^ . While fully automated integration of the scMayoBase annotation results into the pipeline would be possible however some issues are still in favor of manual curation as some cell type definitions are not completely fixed, e.g. in the kidney two distinct datasets for PT and “PT brush border” may reflect the same cell type, intercalated and principal cells may have overlap with distal convoluted tubule and collecting duct cells. Furthermore, in some cases where multiple cell types have similarly high scores the second highest score may be more appropriate, e.g. because the cell type of the highest score appeared multiple times and the second-highest-scored cell type is more specific and known to be contained in the sample. By the advancement of the single-cell technologies existing cell type definitions may be extended, new cell types may be identified which often are sub-types of existing cell types. Concerning this, the fine-tuning of the single cell clustering analysis is crucial: a low resolution will deliver few well-defined clusters but may miss important cell type while a high resolution will deliver also rare cell types and many sub-types of large cell populations which may reflect new sub-cell-types but will not always be reproducible. Thus, it is important to find a good balance between both extremes. In our study, we used a resolution of 0.8 for kidney which enabled detection of low abundant population of podocytes on the one hand and on the other found several sub-clusters of PT cells which might be merged or further explored for sub-types. For brain, a resolution of 0.5 was sufficient for identifying all relevant cell types in only 11 clusters while in kidney in both datasets there were more than 20 clusters. Besides all the obvious advantages of scRNA-seq there should be awareness of the disadvantages: a systemic problem is the lower amount of biological material from the single cells which can be further subjected to stress by the isolation procedure and in consequence there are only a few thousand unique transcripts from a single cell, far less than the one or multiple ten thousands of transcripts from bulk material ^24^ . Thus, in comparison to bulk RNA-seq, scRNA-seq display a higher level of noise and more technical and biological variation^25^ – the latter however as a main topic of investigation. The lower amount of RNA leads to false negatives, here also called “dropouts” ^26^ – genes expressed but not detected. Another issue which can falsify results are dead and low-quality cells. These may computationally be filtered by quality criteria such as thresholds for total numbers of expressed genes per cell or percentages of mitochondrial genes which are supposed to be high in broken cells. However, the proportion of mitochondrial genes is also high in parenchymal kidney cell types such as PT ^15^ so that a well-balanced threshold should be chosen. While we used 50%, Mc Evoy et al. used a threshold of 40% mitochondrial genes and review that others retained cells with a maximum of 25-80% mitochondrial reads ^15^ . Single-cell techniques obviously offer a plethora of novel applications but taking into account the drawbacks listed and also the high variation between cell type distributions in datasets of the same organ we found the question emerges if for conventional gene expression analyses the bulk approach might be superior. Thus, a quality study systematically comparing the accuracy and reproducibility of bulk and single-cell approaches summarized to the bulk level would be of great benefit for researchers to help them decide if for their scientific question the bulk or the single-cell approach is preferable.

Results derived from deconvolution obviously depend on the scRNA-seq dataset used, in particular as a Bayesian method will integrate the underlying a priori distribution of cell types. Thus, while it is clear that at least the scRNA-seq data should be from the same organ as the bulk data, e.g. both datasets from kidney, further restrictions for organ sub-regions, sex or other factors may additionally improve the results.

We conclude, that our study on deconvolution of bulk transcriptome data using scRNA-seq data revealed that most relevant organ-specific and immune cell types could be detected in bulk iPSC-derived kidney and brain organoid transcriptome data. The results depend on the scRNA-seq datasets chosen and on their analysis with appropriate cluster resolution and annotation for which we applied the tools Seurat and scMayoBase integrating a comprehensive set of tissue-specific datasets. The deconvolution via InstaPrism could qualitatively identify the cell types contained in the bulk transcriptome data, also low abundant populations such as podocytes in kidney and microglia in brain. However, the proportions of cell types vary among the scRNA-seq datasets employed and we hypothesize will be most exact when the scRNA-seq dataset most closely resembles the bulk data.

## Supporting information

Supplemental Item 1

## Acknowledgements

James Adjaye acknowledges support from the Medical faculty of the Heinrich-Heine University, Duesseldorf.

## Author contributions

WW analyzed the single-cell and bulk data and wrote the manuscript. JA supervised the work, co-wrote the manuscript and gave the final approval.

## Methods

### Sample collection of kidney and brain scRNA-seq and bulk iPSC-derived kidney and brain organoid transcriptome data

Kidney and brain scRNA-seq datasets used in this study are listed in Table S1 and were downloaded from NCBI GEO (National Center for Biotechnology Information, Gene Expression Omnibus). Bulk kidney and brain organoid gene expression microarray datasets for which the cell type distribution should be estimated were from previous publications of our lab and are available as indicated in Table S1. The bulk data were pre-processed and normalized as described in our previous publications by Martins et al. ^22^, Nguyen et al. ^13^ and Pranty et al. ^21^ . Furthermore, the normalized bulk transcriptome data were summarized to unique gene symbols using the probesets with the maximum expression in the mean of all samples.

### Cluster analysis of kidney and brain-biopsy scRNA-seq data

Kidney and brain scRNA-seq datasets listed in Table S1 were processed in the R / Bioconductor environment (R version 4.3.2) ^27^ . The R package Seurat (Version 5.1.0) ^6^ was used for integration of the scRNA-seq samples, further preprocessing, finding of the clusters and markers associated with the clusters. The code implemented for the single-cell and deconvolution analyses is provided in supplemental information item SI1. For the integration we followed the standard workflow: data was normalized, scaled, variable features were found with the selection method “vst”, PCA was performed, the integration features were selected and the integration anchors were identified with the fast method of reciprocal PCA (‘RPCA’). Based on the identified integration anchors, data was integrated and layers were joined. The integrated data was scaled, PCA and UMAP were performed with 30 dimensions, neighbours and clusters were found with a resolution of 0.8. Markers of all clusters were identified with the function FindAllMarkers parametrized to only test genes that are detected in a minimum fraction of min.pct=0.25 cells and with a threshold for the log-fold-change of 0.25. The markers identified in this way were used as input for the scMayoMap package which is based on a database of scRNA-seq and bulk RNA-seq datasets associated with distinct tissues and can therefore estimate distinct cell type clusters. The function call to scMayoMap was parametrized with the tissue “brain” for the dataset GSE104276 and with the tissue “kidney” for the datasets GSE241302 and GSE202109. ScMayoMap produces a dot plot (and corresponding tables) indicating the cluster-associated cell types with the highest score and in some cases second highest score. Clusters not annotated by scMayoMap were annotated via the EnrichR webtool ^17^ uploading the set of marker genes and using ARCHS4-Tissue datasets in EnrichR.

A lookup-table of cell types associated with clusters based on these results was read and used to rename the cluster ids for the subsequent UMAP plot and calculation of fractions of cell types. Fractions of cell types were calculated by dividing the number of single cells of a distinct cell type by the number of all single cells.

### Deconvolution analysis of iPSC-derived kidney and brain organoid bulk RNA-seq data based on scRNA-seq data as reference

Deconvolution was performed in R (version 4.3.2) / Bioconductor like the Seurat analysis. Biopsy-derived kidney and brain scRNA-seq expression data was reduced to the set of unique marker genes of all clusters. As for the kidney sc-dataset GSE241302 16GB main memory were not sufficient to store the Seurat and deconvolution data in parallel as such the work flow was split in a Seurat R run which wrote the scRNA-seq expression reduced to marker genes onto disk and a second R run reading this data and performing the deconvolution. For the other datasets, 16 GB main memory was sufficient to store Seurat and deconvolution data in parallel. Besides, the sc-data the bulk organoid microarray data – pre-processed and reduced to unique genes as described above - was read. Sc- and bulk expression data were reduced to the inter-section set of genes and merged. Batch effects of the merged data were removed via the method ComBat from the R package sva ^18^ and data was normalized with quantile normalization ^19^ . Deconvolution was performed via the R package InstaPrism ^5^ which is an efficient implementation of BayesPrism ^4^ . The call to InstaPrism is parametrized with the sc-part of the merged data, the bulk part of the merged data and the clusters associated with single cells via Seurat. Results of InstaPrism are the posterior theta values which estimate the proportions of clusters. These are then associated with cell types via the scMayoMap lookup table and summarized in a table and a pie chart.

### Data and code availability

All original codes are available as supplemental information.

