## Supplemental Item 1 for "Single cell data enables dissecting cell types present in bulk transcriptome data": wruck_etal_eval_GSE104276_seurat.R

```
################################################################################################
# R code for analysis of sc-RNAseq developmental brain dataset GSE104276
# and deconvolution of two brain organoid microarray datasets bundled with this software
# R version 4.3.2
#
# input:
# sc-data to be downloaded from NCBI GEO GSE104276: GSE104276_all_pfc_2394_UMI_TPM_NOERCC.txt
# bulk brain organoid microarray dataset1: bilirubin_brainorgs_sym.txt
# bulk brain organoid microarray dataset2: NBS_brainorgs_sym.txt
# celltype cluster lookup table: clusterdesc_GSE104276.txt
#
# author: Wasco Wruck, 2024
# institute: Institute for Stem Cell Research and Regenerative Medicine
# University Clinic Duesseldorf, Germany
################################################################################################
library(Seurat) # R4.3
library(gplots)
library(scMayoMap)
library(ggplot2)
library(svglite)
library(preprocessCore)
library(sva)
library(InstaPrism)
library(plotrix)

# get zhong nature 2018 TPM table from NCBI GEO GSE104276, unzip and open in Excel
# In the 1st column there are genesymbols but a header is missing.
# Therefore, copy all header names starting in cell A1 to B1 and write "genesymbol" to cell A1.
# save as tab-delimitted .txt file
t1 <- read.table("GSE104276_all_pfc_2394_UMI_TPM_NOERCC.txt", header=TRUE, sep="\t",row.names=1,as.is=TRUE)
# remove rows with zero counts
nullRows=which(rowSums(t1)==0)
t1=t1[-(nullRows),]
seurat_R1 <- CreateSeuratObject(counts = t1)

# Run the standard workflow for visualization and clustering
seurat_R1 <- NormalizeData(seurat_R1, normalization.method = "LogNormalize", scale.factor = 10000)
seurat_R1 <- ScaleData(seurat_R1, verbose = FALSE)
seurat_R1 <- FindVariableFeatures(seurat_R1, selection.method = "vst", nfeatures = 2000)
seurat_R1 <- RunPCA(seurat_R1, npcs = 30, verbose = FALSE)
seurat_R1 <- RunUMAP(seurat_R1, reduction = "pca", dims = 1:30)
seurat_R1 <- FindNeighbors(seurat_R1, reduction = "pca", dims = 1:30)
seurat_R1 <- FindClusters(seurat_R1, resolution = 0.5)

DefaultAssay(seurat_R1) <- "RNA"
all.markers <- FindAllMarkers(seurat_R1, only.pos = TRUE, min.pct = 0.25, logfc.threshold = 0.25)
write.table(all.markers,file="markers_GSE104276.txt",sep="\t",quote=F,row.names=F)

#scmayomap: identify celltypes of clusters
scMayoMap.obj <- scMayoMap(data = all.markers, database=scMayoMapDatabase, tissue = 'brain')
plt <- scMayoMap.plot(scMayoMap.object = scMayoMap.obj)
ggsave("scMayoMap_plot_GSE104276.svg",plot=plt,width=10,height=6)
write.table(scMayoMap.obj$res,file="scMayoMap_GSE104276_table.txt",sep="\t",quote=F,row.names=F)
write.table(scMayoMap.obj$markers,file="scMayoMap_GSE104276_markers.txt",sep="\t",quote=F,row.names=F)

# manually assign celltypes (column 2) found by scMayoMap to cluster nos. (col. 1) in Excel and save as tab-del. table
# read cluster associations by scMayoMap (manually assigned celltypes (column 2) to cluster nos. (col. 1)in tab-del. table)
cludesc=read.table("clusterdesc_GSE104276.txt",sep="\t",header=T,row.names=1)
new.cluster.ids <- cludesc[,1]
names(new.cluster.ids) <- levels(seurat_R1)
seurat_R1 <- RenameIdents(seurat_R1, new.cluster.ids)
pdf("umap_clusters_desc_GSE104276.pdf",width=5,height=5)
DimPlot(seurat_R1, reduction = "umap", label = TRUE,label.size=3,pt.size = 0.5) + NoLegend()
dev.off()
ct=unique(all.markers[,"gene"])
m=match(ct,rownames(t1))
t2=t1[m,]
# LogNormalize: Feature counts for each cell are divided by the total counts for that cell and multiplied 
#   by the scale.factor. This is then natural-log transformed using log1p
sums=colSums(t2)
mdsums=median(sums)
for (i in 1:dim(t2)[2]){
  t2[,i]=log(1 + t2[,i] * mdsums / sums[i])
}
# write normalized data and clusters - uncomment if needed
# write.table(t2,file="GSE104276_devbrain_counts_CmarkersU.txt", col.names=TRUE,quote=F, sep="\t")
# write.table(as.character(seurat_R1@meta.data$seurat_clusters),file="GSE104276_devbrain_sctypes.txt",row.names=F,col.names=F,quote=F, sep="\t")
ac=as.character(seurat_R1@meta.data$seurat_clusters)
has_NA=apply(t2,2,function(x){return(sum(is.na(x)) > 0)})# samples containing NAs
t2m=as.matrix(2^t2[,-(which(has_NA))]) # remove NAs from matrix
ac2=ac[-(which(has_NA))] # remove NAs from clusters
# get fractions of celltypes
fractions=table(ac2)/sum(table(ac2))
ct=read.table("clusterdesc_GSE104276.txt", header=T) # read cluster descriptions
names(fractions)=ct[match(names(fractions),ct$cluster),"celltype"]
f=data.frame(fractions)
fs=f[order(f$Freq,decreasing=T),]
write.table(fs,file="fractions_GSE104276.txt",col.names=T,row.names=F,sep="\t",quote=F)

############## DECONVOLUTION ###################################

# 1st BULK brain organoid trancriptome bilirubin_normalized_data
pt2 <- read.table("bilirubin_brainorgs_sym.txt", header=TRUE, sep="\t",row.names=1,as.is=TRUE)
symbol=rownames(pt2) # gene symbol
m=which(symbol %in% rownames(t2m)) # get all matches
cexprs=pt2[m,]
cprobes=rownames(cexprs)
tr_red=t2m[rownames(cexprs),]
mat=as.matrix(cbind(tr_red,cexprs))
mat2=ComBat(dat=mat,batch=c(rep(1,dim(tr_red)[2]),rep(2,dim(cexprs)[2])),prior.plots =FALSE) # remove batch effects
matn=normalize.quantiles(mat2)
tr_red2=matn[,1:dim(tr_red)[2]]
cexprs2=matn[,(dim(tr_red)[2]+1):dim(mat)[2]]
colnames(tr_red2)=colnames(tr_red)
rownames(tr_red2)=rownames(tr_red)
colnames(cexprs2)=colnames(cexprs)
rownames(cexprs2)=rownames(cexprs)

# InstaPrism deconvolution
InstaPrism.res = InstaPrism(input_type = 'raw',sc_Expr = tr_red2,bulk_Expr = cexprs2,
                    cell.type.labels = ac2,cell.state.labels = ac2)
# write prediction matrix
orderedsamples=c("UJ_24h_ctrl","UJ_24h_bilirubin","UJ_72h_ctrl","UJ_72h_bilirubin","C2_24h_ctrl","C2_24h_bilirubin","C2_72h_ctrl","C2_72h_bilirubin")
prediction.matrix=InstaPrism.res@Post.ini.cs@theta[,orderedsamples]
ct=read.table("clusterdesc_GSE104276.txt", header=T) # read cluster descriptions
m=match(rownames(prediction.matrix),ct[,1])
rownames(prediction.matrix)=paste(ct[m,2],rownames(prediction.matrix),sep="_")
write.table(prediction.matrix,file="prediction.matrix_GSE104276_bili.txt",col.names=T,row.names=T,sep="\t",quote=F)

# pie chart
pdt=read.table("prediction.matrix_GSE104276_bili.txt", header=T,sep="\t")
s=c(30,129,348,135,147,358,459,168,483,465,654,522,251,489,440,126,172,555,460,219,498,38,407,302,169,132,329,196,231,486)
slices=prediction.matrix[,grep("UJ_72h_ctrl",colnames(prediction.matrix))]
labels=paste(rownames(prediction.matrix),sprintf(" %.02f",slices*100),"%",sep="")
pdf("pie_ct_GSE104276_UJ_72h_ctrl.pdf")
randcol=colors()[s]
pietitle="cell types UJ_72h_ctrl"
pie3D(slices,labels=labels,explode=0.1,main=pietitle,col=randcol,labelcex=0.6,mar=c(8,4,4,4))
dev.off()
f=data.frame(celltype=rownames(prediction.matrix),percentage=slices*100)
f=f[order(f$percentage,decreasing=TRUE),]
write.table(f,file="percentages_GSE104276_UJ_72h_ctrl.txt",col.names=T,row.names=F,sep="\t",quote=F)

# 2nd BULK brain organoids (Nijmegen-Breakage syndrome - BS) trancriptome NBS_normalized_data
pt2 <- read.table("NBS_brainorgs_sym.txt", header=TRUE, sep="\t",row.names=1,as.is=TRUE)
symbol=rownames(pt2) # gene symbol
m=which(symbol %in% rownames(t2m)) # get all matches
cexprs=pt2[m,]
cprobes=rownames(cexprs)
tr_red=t2m[rownames(cexprs),]
mat=as.matrix(cbind(tr_red,cexprs))
mat2=ComBat(dat=mat,batch=c(rep(1,dim(tr_red)[2]),rep(2,dim(cexprs)[2])),prior.plots =FALSE) # remove batch effects
matn=normalize.quantiles(mat2)
tr_red2=matn[,1:dim(tr_red)[2]]
cexprs2=matn[,(dim(tr_red)[2]+1):dim(mat)[2]]
colnames(tr_red2)=colnames(tr_red)
rownames(tr_red2)=rownames(tr_red)
colnames(cexprs2)=colnames(cexprs)
rownames(cexprs2)=rownames(cexprs)

# InstaPrism deconvolution
InstaPrism.res = InstaPrism(input_type = 'raw',sc_Expr = tr_red2,bulk_Expr = cexprs2,
                    cell.type.labels = ac2,cell.state.labels = ac2)
# write prediction matrix
orderedsamples=c("A4_20","UJ_20","NBS1_20","NBS8_20","A4_40","UJ_40","UJ_40Bleo","NBS1_40","NBS1_40_2","NBS1_40Bleo","NBS1_40Bleo_2")
prediction.matrix=InstaPrism.res@Post.ini.cs@theta[,orderedsamples]
ct=read.table("clusterdesc_GSE104276.txt", header=T) # read cluster descriptions
m=match(rownames(prediction.matrix),ct[,1])
rownames(prediction.matrix)=paste(ct[m,2],rownames(prediction.matrix),sep="_")
write.table(prediction.matrix,file="prediction.matrix_GSE104276_NBS.txt",col.names=T,row.names=T,sep="\t",quote=F)

# pie chart
pdt=read.table("prediction.matrix_GSE104276_NBS.txt", header=T,sep="\t")
s=c(30,129,348,135,147,358,459,168,483,465,654,522,251,489,440,126,172,555,460,219,498,38,407,302,169,132,329,196,231,486)
slices=prediction.matrix[,which(colnames(prediction.matrix)=="UJ_40")]
labels=paste(rownames(prediction.matrix),sprintf(" %.02f",slices*100),"%",sep="")
pdf("pie_ct_GSE104276_UJ_40.pdf")
randcol=colors()[s]
pietitle="cell types UJ_40"
pie3D(slices,labels=labels,explode=0.1,main=pietitle,col=randcol,labelcex=0.6,mar=c(8,4,4,4))
dev.off()
f=data.frame(celltype=rownames(prediction.matrix),percentage=slices*100)
f=f[order(f$percentage,decreasing=TRUE),]
write.table(f,file="percentages_GSE104276_UJ_40.txt",col.names=T,row.names=F,sep="\t",quote=F)
```
