## Supplemental Item 1 for "Single cell data enables dissecting cell types present in bulk transcriptome data": wruck_etal_eval_GSE202109_seurat.R

```
################################################################################################
# R code for analysis of sc-RNAseq kidney dataset GSE202109
# and deconvolution of one kidney organoid microarray datasets bundled with this software
# R version 4.3.2
#
# input:
# sc-data to be downloaded from NCBI GEO GSE202109: 
# matrix, barcode and features .gz files of all samples (except the CD45) from NCBI GEO GSE202109:
# matrix files: 
# GSM6094655_HKB14_matrix.mtx.gz  GSM6094658_HKB17_matrix.mtx.gz  GSM6094668_HKB32_matrix.mtx.gz
# GSM6094656_HKB15_matrix.mtx.gz  GSM6094659_HKB18_matrix.mtx.gz  GSM6094669_HKB33_matrix.mtx.gz
# GSM6094657_HKB16_matrix.mtx.gz  GSM6094667_HKB31_matrix.mtx.gz  GSM6094670_HKB34_matrix.mtx.gz
# and corresponding barcodes and features files
# bulk kidney organoid microarray dataset: PAN_kidneyorgs_sym.txt
# celltype cluster lookup table: clusterdesc_GSE202109.txt
#
# author: Wasco Wruck, 2024
# institute: Institute for Stem Cell Research and Regenerative Medicine
# University Clinic Duesseldorf, Germany
################################################################################################
library(Seurat) # R4.3.2
library(gplots)
library(scMayoMap)
library(ggplot2)
library(svglite)
library(preprocessCore)
library(sva)
library(InstaPrism)
library(plotrix)

# get matrix, barcode and features .gz files of all samples (except the CD45) from NCBI GEO GSE202109:
# matrix files: 
# GSM6094655_HKB14_matrix.mtx.gz  GSM6094658_HKB17_matrix.mtx.gz  GSM6094668_HKB32_matrix.mtx.gz
# GSM6094656_HKB15_matrix.mtx.gz  GSM6094659_HKB18_matrix.mtx.gz  GSM6094669_HKB33_matrix.mtx.gz
# GSM6094657_HKB16_matrix.mtx.gz  GSM6094667_HKB31_matrix.mtx.gz  GSM6094670_HKB34_matrix.mtx.gz
# and corresponding barcodes and features files
expression_matrix <- ReadMtx( # M35
  mtx = "GSM6094655_HKB14_matrix.mtx.gz", features = "GSM6094655_HKB14_features.tsv.gz",
  cells = "GSM6094655_HKB14_barcodes.tsv.gz"
)
seurat_R1 <- CreateSeuratObject(counts = expression_matrix)
DefaultAssay(seurat_R1) <- "RNA"
seurat_R1[["percent.mt"]] <- PercentageFeatureSet(seurat_R1, pattern = "^MT-")
seurat_R1 <- subset(seurat_R1, subset = nFeature_RNA > 200 & percent.mt < 50)

expression_matrix <- ReadMtx( # M65
  mtx = "GSM6094656_HKB15_matrix.mtx.gz", features = "GSM6094656_HKB15_features.tsv.gz",
  cells = "GSM6094656_HKB15_barcodes.tsv.gz"
)
seurat_R2 <- CreateSeuratObject(counts = expression_matrix)
DefaultAssay(seurat_R2) <- "RNA"
seurat_R2[["percent.mt"]] <- PercentageFeatureSet(seurat_R2, pattern = "^MT-")
seurat_R2 <- subset(seurat_R2, subset = nFeature_RNA > 200 & percent.mt < 50)

#
expression_matrix <- ReadMtx(
  mtx = "GSM6094657_HKB16_matrix.mtx.gz", features = "GSM6094657_HKB16_features.tsv.gz",
  cells = "GSM6094657_HKB16_barcodes.tsv.gz"
)
seurat_R3 <- CreateSeuratObject(counts = expression_matrix)
DefaultAssay(seurat_R3) <- "RNA"
seurat_R3[["percent.mt"]] <- PercentageFeatureSet(seurat_R3, pattern = "^MT-")
seurat_R3 <- subset(seurat_R3, subset = nFeature_RNA > 200 & percent.mt < 50)

#
expression_matrix <- ReadMtx(
  mtx = "GSM6094658_HKB17_matrix.mtx.gz", features = "GSM6094658_HKB17_features.tsv.gz",
  cells = "GSM6094658_HKB17_barcodes.tsv.gz"
)
seurat_R4 <- CreateSeuratObject(counts = expression_matrix)
DefaultAssay(seurat_R4) <- "RNA"
seurat_R4[["percent.mt"]] <- PercentageFeatureSet(seurat_R4, pattern = "^MT-")
seurat_R4 <- subset(seurat_R4, subset = nFeature_RNA > 200 & percent.mt < 50)

expression_matrix <- ReadMtx( # F64
  mtx = "GSM6094659_HKB18_matrix.mtx.gz", features = "GSM6094659_HKB18_features.tsv.gz",
  cells = "GSM6094659_HKB18_barcodes.tsv.gz"
)
seurat_R5 <- CreateSeuratObject(counts = expression_matrix)
DefaultAssay(seurat_R5) <- "RNA"
seurat_R5[["percent.mt"]] <- PercentageFeatureSet(seurat_R5, pattern = "^MT-")
seurat_R5 <- subset(seurat_R5, subset = nFeature_RNA > 200 & percent.mt < 50)

expression_matrix <- ReadMtx( # F50
  mtx = "GSM6094667_HKB31_matrix.mtx.gz", features = "GSM6094667_HKB31_features.tsv.gz",
  cells = "GSM6094667_HKB31_barcodes.tsv.gz"
)
seurat_R6 <- CreateSeuratObject(counts = expression_matrix)
DefaultAssay(seurat_R6) <- "RNA"
seurat_R6[["percent.mt"]] <- PercentageFeatureSet(seurat_R6, pattern = "^MT-")
seurat_R6 <- subset(seurat_R6, subset = nFeature_RNA > 200 & percent.mt < 50)

#
expression_matrix <- ReadMtx(
  mtx = "GSM6094668_HKB32_matrix.mtx.gz", features = "GSM6094668_HKB32_features.tsv.gz",
  cells = "GSM6094668_HKB32_barcodes.tsv.gz"
)
seurat_R7 <- CreateSeuratObject(counts = expression_matrix)
DefaultAssay(seurat_R7) <- "RNA"
seurat_R7[["percent.mt"]] <- PercentageFeatureSet(seurat_R7, pattern = "^MT-")
seurat_R7 <- subset(seurat_R7, subset = nFeature_RNA > 200 & percent.mt < 50)

#
expression_matrix <- ReadMtx(
  mtx = "GSM6094669_HKB33_matrix.mtx.gz", features = "GSM6094669_HKB33_features.tsv.gz",
  cells = "GSM6094669_HKB33_barcodes.tsv.gz"
)
seurat_R8 <- CreateSeuratObject(counts = expression_matrix)
DefaultAssay(seurat_R8) <- "RNA"
seurat_R8[["percent.mt"]] <- PercentageFeatureSet(seurat_R8, pattern = "^MT-")
seurat_R8 <- subset(seurat_R8, subset = nFeature_RNA > 200 & percent.mt < 50)

#
expression_matrix <- ReadMtx(
  mtx = "GSM6094670_HKB34_matrix.mtx.gz", features = "GSM6094670_HKB34_features.tsv.gz",
  cells = "GSM6094670_HKB34_barcodes.tsv.gz"
)
seurat_R9 <- CreateSeuratObject(counts = expression_matrix)
DefaultAssay(seurat_R9) <- "RNA"
seurat_R9[["percent.mt"]] <- PercentageFeatureSet(seurat_R9, pattern = "^MT-")
seurat_R9 <- subset(seurat_R9, subset = nFeature_RNA > 200 & percent.mt < 50)

slist=list(seurat_R1,seurat_R2,seurat_R3,seurat_R4,seurat_R5,seurat_R6,seurat_R7,seurat_R8,seurat_R9)
# normalize and identify variable features for each dataset independently
slist <- lapply(X = slist, FUN = function(x) {
    x <- NormalizeData(x)
    x <- FindVariableFeatures(x, selection.method = "vst", nfeatures = 2000)
    x <- ScaleData(x,verbose = FALSE)
    x <- RunPCA(x, npcs = 30, verbose = FALSE)
})
# select features that are repeatedly variable across datasets for integration
features <- SelectIntegrationFeatures(object.list = slist)

# faster version with rpca
s.anchors <- FindIntegrationAnchors(object.list = slist,reduction="rpca", anchor.features = features)

# create an 'integrated' data assay
s.combined <- IntegrateData(anchorset = s.anchors)
s.combined[["RNA"]] <- JoinLayers(s.combined[["RNA"]])
#DefaultAssay(s.combined) <- "integrated"

# Run the standard workflow for visualization and clustering
s.combined <- ScaleData(s.combined, verbose = FALSE)
s.combined <- RunPCA(s.combined, npcs = 30, verbose = FALSE)
s.combined <- RunUMAP(s.combined, reduction = "pca", dims = 1:30)
s.combined <- FindNeighbors(s.combined, reduction = "pca", dims = 1:30)
s.combined <- FindClusters(s.combined, resolution = 0.8)

# switch back to the original data
DefaultAssay(s.combined) <- "RNA"
all.markers <- FindAllMarkers(s.combined, only.pos = TRUE, min.pct = 0.25, logfc.threshold = 0.25)
write.table(all.markers,file="markers_GSE202109.txt",sep="\t",quote=F,row.names=F)

# QUICK START (huge file but speeds up if saved before with save.image("GSE202109.RData")
# load("GSE202109.RData")

#scmayomap: identify celltypes of clusters
scMayoMap.obj <- scMayoMap(data = all.markers, database=scMayoMapDatabase, tissue = 'kidney')
plt <- scMayoMap.plot(scMayoMap.object = scMayoMap.obj)
ggsave("scMayoMap_plot_GSE202109.svg",plot=plt,width=12,height=6)
write.table(scMayoMap.obj$res,file="scMayoMap_GSE202109_table.txt",sep="\t",quote=F,row.names=F)
write.table(scMayoMap.obj$markers,file="scMayoMap_GSE202109_markers.txt",sep="\t",quote=F,row.names=F)

# manually assign celltypes (column 2) found by scMayoMap to cluster nos. (col. 1) in Excel and save as tab-del. table
# read cluster associations by scMayoMap (manually assigned celltypes (column 2) to cluster nos. (col. 1)in tab-del. table)
cludesc=read.table("clusterdesc_GSE202109.txt",sep="\t",header=T,row.names=1)
new.cluster.ids <- cludesc[,1]
names(new.cluster.ids) <- levels(s.combined)
s.combined <- RenameIdents(s.combined, new.cluster.ids)
pdf("umap_clusters_desc_GSE202109.pdf",width=5,height=5)
DimPlot(s.combined, reduction = "umap", label = TRUE,label.size=3,pt.size = 0.5) + NoLegend()
dev.off()
# write counts reduced to unique cluster markers for test of bulk cell type composition (InstaPrism)
marku=unique(all.markers[,"gene"])
t2=s.combined[["RNA"]]$data[marku,]
# write normalized data and clusters, takes much time and disk space! - uncomment if needed
# write.table(x,file="GSE202109_counts_CmarkersU.txt", col.names=TRUE,quote=F, sep="\t")
# write.table(s.combined@meta.data$seurat_clusters,file="GSE202109_clusters.txt",col.names=F,row.names=F,quote=F,sep="\t")
ac=as.character(s.combined@meta.data$seurat_clusters)
t2m=as.matrix(2^t2) 
# get fractions of celltypes
fractions=table(ac)/sum(table(ac))
ct=read.table("clusterdesc_GSE202109.txt", header=T) # read cluster descriptions
names(fractions)=ct[match(names(fractions),ct$cluster),"celltype"]
f=data.frame(fractions)
fs=f[order(f$Freq,decreasing=T),]
write.table(fs,file="fractions_GSE202109.txt",col.names=T,row.names=F,sep="\t",quote=F)

############## DECONVOLUTION ###################################

# 1st BULK brain organoid trancriptome bilirubin_normalized_data
pt2 <- read.table("PAN_kidneyorgs_sym.txt", header=TRUE, sep="\t",row.names=1,as.is=TRUE)
symbol=rownames(pt2) # gene symbol
m=which(symbol %in% rownames(t2m)) # get all matches
cexprs=pt2[m,]
cprobes=rownames(cexprs)
tr_red=t2m[rownames(cexprs),]
mat=as.matrix(cbind(tr_red,cexprs))
mat2=ComBat(dat=mat,batch=c(rep(1,dim(tr_red)[2]),rep(2,dim(cexprs)[2])),prior.plots =FALSE) # remove batch effects
matn=normalize.quantiles(mat2)
tr_red2=matn[,1:dim(tr_red)[2]]
cexprs2=matn[,(dim(tr_red)[2]+1):dim(mat)[2]]
colnames(tr_red2)=colnames(tr_red)
rownames(tr_red2)=rownames(tr_red)
colnames(cexprs2)=colnames(cexprs)
rownames(cexprs2)=rownames(cexprs)

# InstaPrism deconvolution
InstaPrism.res = InstaPrism(input_type = 'raw',sc_Expr = tr_red2,bulk_Expr = cexprs2,
                    cell.type.labels = ac,cell.state.labels = ac)
# write prediction matrix
prediction.matrix=InstaPrism.res@Post.ini.cs@theta
ct=read.table("clusterdesc_GSE202109.txt", header=T) # read cluster descriptions
m=match(rownames(prediction.matrix),ct[,1])
rownames(prediction.matrix)=paste(ct[m,2],rownames(prediction.matrix),sep="_")
write.table(prediction.matrix,file="prediction.matrix_GSE202109_PAN.txt",col.names=T,row.names=T,sep="\t",quote=F)

# pie chart
pdt=read.table("prediction.matrix_GSE202109_PAN.txt", header=T,sep="\t")
s=c(30,129,348,135,147,358,459,168,483,465,654,522,251,489,440,126,172,555,460,219,498,38,407,302,169,132,329,196,231,486)
slices=prediction.matrix[,grep("UMK1_con_1",colnames(prediction.matrix))]
labels=paste(rownames(prediction.matrix),sprintf(" %.02f",slices*100),"%",sep="")
pdf("pie_ct_GSE202109_UMK1_con_1.pdf")
randcol=colors()[s]
pietitle="cell types UMK1_con_1"
pie3D(slices,labels=labels,explode=0.1,main=pietitle,col=randcol,labelcex=0.6,mar=c(8,4,4,4))
dev.off()
f=data.frame(celltype=rownames(prediction.matrix),percentage=slices*100)
f=f[order(f$percentage,decreasing=TRUE),]
write.table(f,file="percentages_GSE202109_UMK1_con_1.txt",col.names=T,row.names=F,sep="\t",quote=F)
```
