## Supplemental Item 1 for "Single cell data enables dissecting cell types present in bulk transcriptome data": wruck_etal_eval_GSE241302_deconv.R

```
################################################################################################
# R code for deconvolution based on sc-RNAseq kidney dataset GSE241302
# with one kidney organoid microarray datasets bundled with this software
# (2nd part, first run seurat script)
# R version 4.3.2
#
# input:
# sc-data written by seurat script: 
# sc-expression data reduced to markers: GSE241302_counts_CmarkersU.txt
# clusters: GSE241302_clusters.txt
# bulk kidney organoid microarray dataset: PAN_kidneyorgs_sym.txt
# celltype cluster lookup table: clusterdesc_GSE241302.txt
#
# author: Wasco Wruck, 2024
# institute: Institute for Stem Cell Research and Regenerative Medicine
# University Clinic Duesseldorf, Germany
################################################################################################
library(gplots)
library(InstaPrism) # R4.3.2
library(plotrix)
library(Biobase)
library(preprocessCore)
library(sva)
library(plotrix)

############## DECONVOLUTION ###################################
# on a core i7 laptop with 16GB RAM R4.3.2 crashed when doing Seurat and InstaPrism
# in one run. Therefore, Seurat results are written as count table of reduced markers and
# clusters and read here.
# read GSE241302 count table reduced to unique cluster markers directly 
t2 <- read.table("GSE241302_counts_CmarkersU.txt", header=TRUE, sep="\t",row.names=1,as.is=TRUE)
tkm <- read.table("GSE241302_clusters.txt", header=F, sep="\t") # read clusters from file
t2m=as.matrix(2^t2)
ac=as.character(tkm[,1])

# 1st BULK kidney organoid trancriptome PAN_kidneyorgs_normalized_data
pt2 <- read.table("PAN_kidneyorgs_sym.txt", header=TRUE, sep="\t",row.names=1,as.is=TRUE)
symbol=rownames(pt2) # gene symbol
m=which(symbol %in% rownames(t2m)) # get all matches
cexprs=pt2[m,]
cprobes=rownames(cexprs)
tr_red=t2m[rownames(cexprs),]
mat=as.matrix(cbind(tr_red,cexprs))
mat2=ComBat(dat=mat,batch=c(rep(1,dim(tr_red)[2]),rep(2,dim(cexprs)[2])),prior.plots =FALSE) # remove batch effects
matn=normalize.quantiles(mat2)
tr_red2=matn[,1:dim(tr_red)[2]]
cexprs2=matn[,(dim(tr_red)[2]+1):dim(mat)[2]]
colnames(tr_red2)=colnames(tr_red)
rownames(tr_red2)=rownames(tr_red)
colnames(cexprs2)=colnames(cexprs)
rownames(cexprs2)=rownames(cexprs)

# InstaPrism deconvolution
InstaPrism.res = InstaPrism(input_type = 'raw',sc_Expr = tr_red2,bulk_Expr = cexprs2,
                    cell.type.labels = ac,cell.state.labels = ac)
# write prediction matrix
prediction.matrix=InstaPrism.res@Post.ini.cs@theta
ct=read.table("clusterdesc_GSE241302.txt", header=T) # read cluster descriptions
m=match(rownames(prediction.matrix),ct[,1])
rownames(prediction.matrix)=paste(ct[m,2],rownames(prediction.matrix),sep="_")
write.table(prediction.matrix,file="prediction.matrix_GSE241302_PAN.txt",col.names=T,row.names=T,sep="\t",quote=F)

# pie chart
pdt=read.table("prediction.matrix_GSE241302_PAN.txt", header=T,sep="\t")
s=c(30,129,348,135,147,358,459,168,483,465,654,522,251,489,440,126,172,555,460,219,498,38,407,302,169,132,329,196,231,486)
slices=prediction.matrix[,grep("UMK1_con_1",colnames(prediction.matrix))]
labels=paste(rownames(prediction.matrix),sprintf(" %.02f",slices*100),"%",sep="")
pdf("pie_ct_GSE241302_UMK1_con_1.pdf")
randcol=colors()[s]
pietitle="cell types UMK1_con_1"
pie3D(slices,labels=labels,explode=0.1,main=pietitle,col=randcol,labelcex=0.6,mar=c(8,4,4,4))
dev.off()
f=data.frame(celltype=rownames(prediction.matrix),percentage=slices*100)
f=f[order(f$percentage,decreasing=TRUE),]
write.table(f,file="percentages_GSE241302_UMK1_con_1.txt",col.names=T,row.names=F,sep="\t",quote=F)
```
