## Supplemental Item 1 for "Single cell data enables dissecting cell types present in bulk transcriptome data": wruck_etal_eval_GSE241302_seurat.R

```
################################################################################################
# R code for analysis of sc-RNAseq kidney dataset GSE241302
# and deconvolution of one kidney organoid microarray datasets bundled with this software
# R version 4.3.2
#
# input:
# sc-data to be downloaded from NCBI GEO GSE241302: 
# matrix, barcode and features .gz files of all samples (except the CD45) from NCBI GEO GSE241302:
# matrix files: 
# GSM7720822_R1_MN_matrix.mtx.gz  GSM7720823_R2_MN_matrix.mtx.gz  GSM7720824_R3_MN_matrix.mtx.gz
# GSM7720825_R1_NC_matrix.mtx.gz 
# and corresponding barcodes and features files
# celltype cluster lookup table: clusterdesc_GSE241302.txt
# output:
# sc-expression data reduced to markers: GSE241302_counts_CmarkersU.txt
# clusters: GSE241302_clusters.txt
#
# author: Wasco Wruck, 2024
# institute: Institute for Stem Cell Research and Regenerative Medicine
# University Clinic Duesseldorf, Germany
################################################################################################
library(Seurat) # R4.3.2
library(gplots)
library(scMayoMap)
library(ggplot2)
library(svglite)
library(preprocessCore)
library(sva)
library(InstaPrism)
library(plotrix)

# get matrix, barcode and features .gz files of following samples from NCBI GEO GSE241302:
# matrix files: 
# GSM7720822_R1_MN_matrix.mtx.gz  GSM7720823_R2_MN_matrix.mtx.gz  GSM7720824_R3_MN_matrix.mtx.gz
# GSM7720825_R1_NC_matrix.mtx.gz 
# and corresponding barcodes and features files
expression_matrix <- ReadMtx(
  mtx = "GSM7720822_R1_MN_matrix.mtx.gz", features = "GSM7720822_R1_MN_features.tsv.gz",
  cells = "GSM7720822_R1_MN_barcodes.tsv.gz"
)
seurat_R1_MN <- CreateSeuratObject(counts = expression_matrix)

expression_matrix <- ReadMtx(
  mtx = "GSM7720823_R2_MN_matrix.mtx.gz", features = "GSM7720823_R2_MN_features.tsv.gz",
  cells = "GSM7720823_R2_MN_barcodes.tsv.gz"
)
seurat_R2_MN <- CreateSeuratObject(counts = expression_matrix)

expression_matrix <- ReadMtx(
  mtx = "GSM7720824_R3_MN_matrix.mtx.gz", features = "GSM7720824_R3_MN_features.tsv.gz",
  cells = "GSM7720824_R3_MN_barcodes.tsv.gz"
)
seurat_R3_MN <- CreateSeuratObject(counts = expression_matrix)

expression_matrix <- ReadMtx(
  mtx = "GSM7720825_R1_NC_matrix.mtx.gz", features = "GSM7720825_R1_NC_features.tsv.gz",
  cells = "GSM7720825_R1_NC_barcodes.tsv.gz"
)
seurat_R1_NC <- CreateSeuratObject(counts = expression_matrix)

slist=list(seurat_R1_MN,seurat_R2_MN,seurat_R3_MN,seurat_R1_NC)
# normalize and identify variable features for each dataset independently
slist <- lapply(X = slist, FUN = function(x) {
    x <- NormalizeData(x)
    x <- FindVariableFeatures(x, selection.method = "vst", nfeatures = 2000)
    x <- ScaleData(x,verbose = FALSE)
    x <- RunPCA(x, npcs = 30, verbose = FALSE)
})
# select features that are repeatedly variable across datasets for integration
features <- SelectIntegrationFeatures(object.list = slist)

# faster version with rpca
s.anchors <- FindIntegrationAnchors(object.list = slist,reduction="rpca", anchor.features = features)

# create an 'integrated' data assay
s.combined <- IntegrateData(anchorset = s.anchors)
s.combined[["RNA"]] <- JoinLayers(s.combined[["RNA"]])
#DefaultAssay(s.combined) <- "integrated"

# Run the standard workflow for visualization and clustering
s.combined <- ScaleData(s.combined, verbose = FALSE)
s.combined <- RunPCA(s.combined, npcs = 30, verbose = FALSE)
s.combined <- RunUMAP(s.combined, reduction = "pca", dims = 1:30)
s.combined <- FindNeighbors(s.combined, reduction = "pca", dims = 1:30)
s.combined <- FindClusters(s.combined, resolution = 0.8)

# switch back to the original data
DefaultAssay(s.combined) <- "RNA"
all.markers <- FindAllMarkers(s.combined, only.pos = TRUE, min.pct = 0.25, logfc.threshold = 0.25)
write.table(all.markers,file="markers_GSE241302.txt",sep="\t",quote=F,row.names=F)

# QUICK START (huge file but speeds up if saved before with save.image("GSE241302.RData")
# load("GSE241302.RData")

#scmayomap: identify celltypes of clusters
scMayoMap.obj <- scMayoMap(data = all.markers, database=scMayoMapDatabase, tissue = 'kidney')
plt <- scMayoMap.plot(scMayoMap.object = scMayoMap.obj)
ggsave("scMayoMap_plot_GSE241302.svg",plot=plt,width=12,height=6)
write.table(scMayoMap.obj$res,file="scMayoMap_GSE241302_table.txt",sep="\t",quote=F,row.names=F)
write.table(scMayoMap.obj$markers,file="scMayoMap_GSE241302_markers.txt",sep="\t",quote=F,row.names=F)

# manually assign celltypes (column 2) found by scMayoMap to cluster nos. (col. 1) in Excel and save as tab-del. table
# read cluster associations by scMayoMap (manually assigned celltypes (column 2) to cluster nos. (col. 1)in tab-del. table)
cludesc=read.table("clusterdesc_GSE241302.txt",sep="\t",header=T,row.names=1)
new.cluster.ids <- cludesc[,1]
names(new.cluster.ids) <- levels(s.combined)
s.combined <- RenameIdents(s.combined, new.cluster.ids)
pdf("umap_clusters_desc_GSE241302.pdf",width=5,height=5)
DimPlot(s.combined, reduction = "umap", label = TRUE,label.size=3,pt.size = 0.5) + NoLegend()
dev.off()
# write counts reduced to unique cluster markers for test of bulk cell type composition (InstaPrism)
marku=unique(all.markers[,"gene"])
t2=s.combined[["RNA"]]$data[marku,]
# write normalized data and clusters, takes much time and disk space! 
# needed because Seurat and InstaPrism in one run crashes on a 16GB core i7 laptop
write.table(t2,file="GSE241302_counts_CmarkersU.txt", col.names=TRUE,quote=F, sep="\t")
write.table(s.combined@meta.data$seurat_clusters,file="GSE241302_clusters.txt",col.names=F,row.names=F,quote=F,sep="\t")
ac=as.character(s.combined@meta.data$seurat_clusters)
t2m=as.matrix(2^t2) 
# get fractions of celltypes
fractions=table(ac)/sum(table(ac))
ct=read.table("clusterdesc_GSE241302.txt", header=T) # read cluster descriptions
names(fractions)=ct[match(names(fractions),ct$cluster),"celltype"]
f=data.frame(fractions)
fs=f[order(f$Freq,decreasing=T),]
write.table(fs,file="fractions_GSE241302.txt",col.names=T,row.names=F,sep="\t",quote=F)

############## DECONVOLUTION ###################################
```
